# c-di-AMP inactivates a K^+^/H^+^ antiporter in *Bacillus subtilis*

**DOI:** 10.64898/2026.03.23.713699

**Authors:** Inês Rosa Figueiredo-Costa, Marta Maria Lorga-Gomes, Sara Capela Sousa-Moreira, Isabel Maria Matas, João Henrique Morais-Cabral

**Affiliations:** i3S - Instituto de Investigação e Inovação em Saúde, Universidade do Porto; IBMC – Instituto de Biologia Molecular e Celular, Universidade do Porto

**Keywords:** Potassium transport, Membrane protein, Cyclic diadenosine monophosphate (c-di-AMP), Crystal structure, Structure-function

## Abstract

c-di-AMP is a bacterial second messenger with the crucial role of regulating turgor and osmotic adaptation. Due to the importance of intracellular K^+^ for osmotic balance, c-di-AMP controls the import and export of K^+^ by regulating the activity and transcription level of K^+^ transporters and channels. It has been postulated that c-di-AMP inactivates K^+^ import and activates K^+^ export. To gain a full understanding of the properties the K^+^ machinery in the Gram-positive model organism *Bacillus subtilis* and in particular, of how the machinery is regulated by c-di-AMP, we characterized the molecular properties of CpaA, a cation/H^+^ antiporter that has been shown to bind the dinucleotide. We determined the crystal structure of the cytosolic RCK domain with and without c-di-AMP and performed a functional characterization of full-length CpaA using a fluorescence-based flux assay. We found that c-di-AMP binds on the interface of the RCK-C subdomain but only small structural differences are detected between the apo- and holo-structure. We determined that CpaA is more active at high pH and that it slightly favors K^+^ over Na^+^ for exchange with H^+^. Unexpectedly, CpaA is inactivated by c-di-AMP with a K_1/2_ of inactivation around 1 µM. Our results reinforce the emerging view that regulation of the bacterial K^+^ machinery by c-di-AMP is more complex than previously thought and that a detailed characterization of the molecular properties of the individual protein components and of how their activity is integrated is necessary for a complete view of the machinery physiological function.

## Introduction

c-di-AMP is a bacterial second messenger that is present in many bacterial species, including many pathogens (1). Its primary role is to regulate the protein levels and activity of the proteins that control osmotic adaptation and maintain turgor pressure and volume (1-5). Accordingly, c-di-AMP controls the activity of potassium transporters and channels as K^+^ is central to osmotic regulation in bacteria (3,6-13).

The K^+^ machinery (the combination of K^+^ transporters and channels and their regulatory proteins) of *Bacillus subtilis* has been well characterized. The extensive characterization of the K^+^/H^+^ symporter KimA (12,14), of KtrD and KtrB cation channels and their regulatory proteins KtrA and KtrC (13-17) and of the KhtTU K^+^/H^+^ antiporter (11,14,18,19) has provided a unique global and protein-specific view of the K^+^ machinery in *Bacillus subtilis*, making this model organism a reference system for the study of the central role of K^+^ import and export in bacterial physiology. These studies have clearly established the crucial role of c-di-AMP in the control of the *B. subtilis* K^+^ machinery with the dinucleotide inactivating KimA and the Ktr channels and activating KhtTU. Together with data from other bacterial species (10,20,21), this has resulted in the general proposal that c-di-AMP regulates intracellular K^+^ levels by inactivating K^+^ import and activating K^+^ export (1).

CpaA (also known as YjbQ) is a membrane protein in *B. subtilis* that has been shown to bind c-di-AMP (14). The CpaA polypeptide includes an N-terminal membrane buried domain that belongs to the Cation/Proton Antiporter (CPA) superfamily and as such is thought to assemble as a dimer. In addition, the polypeptide includes a cytosolic Regulating Conductance of K^+^ (RCK) domain at the C terminus. As in other RCK domains, the c-di-AMP binding site is expected to reside at the dimeric interface of the RCK-C subdomain (11,22,23). A study of a CpaA orthologue from *Streptococcus aureus*, sharing 52% identity, revealed the structure of RCK-C with bound c-di-AMP and proposed that this dinucleotide regulates ion transport (2,23).

To gain a full understanding of the properties of the K^+^ machinery in *B. subtilis*, we performed a molecular characterization of CpaA. Unexpectedly, we have determined that this K^+^/H^+^ antiporter, which is expected to function as a K^+^ exporter, is inhibited by c-di-AMP, revealing that regulation of the bacterial K^+^ machinery by this second messenger is more complex than has been thought.

## Results

### Structure of the RCK domain of CpaA with and without c-di-AMP

We first confirmed that the RCK domain of *B. subtilis* CpaA (CpaA-RCK, residues R394 to the C-terminus) was able to bind c-di-AMP using a thermal shift assay with purified protein (Figure 1 and S1). In this assay we also included other compounds, such as pApA (the degradation product of c-di-AMP), c-di-GMP (a related dinucleotide), and Ca^2+^, ATP and ADP previously shown to interact with other RCK proteins. While the majority of these compounds could only slightly change the thermal profile of CpaA-RCK (ΔTm < 2 °C), c-di-AMP increased the melting temperature (Tm) of the protein by more than 6°C (Table S1), strongly indicating that c-di-AMP contributes for CpaA-RCK thermal stability by binding to it.

**Figure 1.**
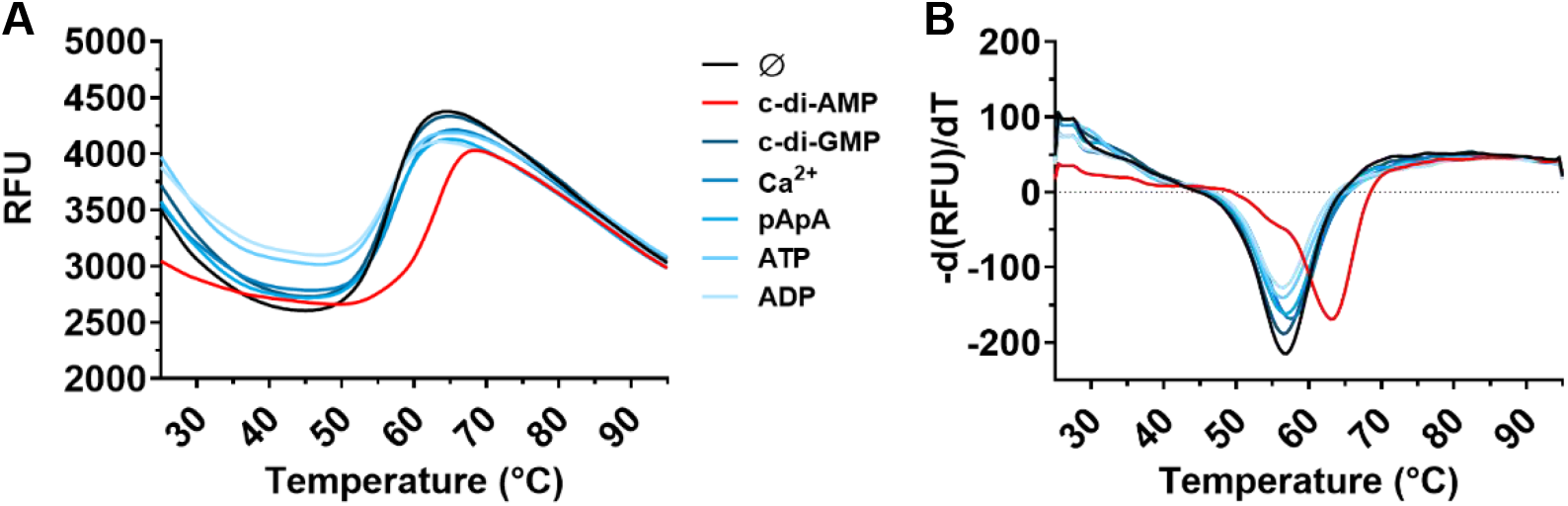
RCK domain of CpaA binds c-di-AMP. **A)** Representative fluorescence curve of Sypro Orange as a function of temperature obtained for the isolated RCK domain of CpaAin the absence and presence of different compounds. **B)** First derivative of the curves in A. Dinucleotides were at 100 µM, while Ca^2+^, ATP and ADP were at 1 mM.

To clarify the molecular interactions established between c-di-AMP and the protein, as well as, to determine the conformational impact of the ligand in the overall structure of the protein, we solved the crystal structure of CpaA-RCK in the absence and presence of the dinucleotide. The structure of the apo-protein at 2.2 Å reveals a typical RCK domain dimer (Figure 2A and Table S2), where each subunit is composed of an N-subdomain (RCK-N), that includes a domain swap-helix from the neighboring dimer subunit, and a smaller C-subdomain (RCK-C).

**Figure 2.**
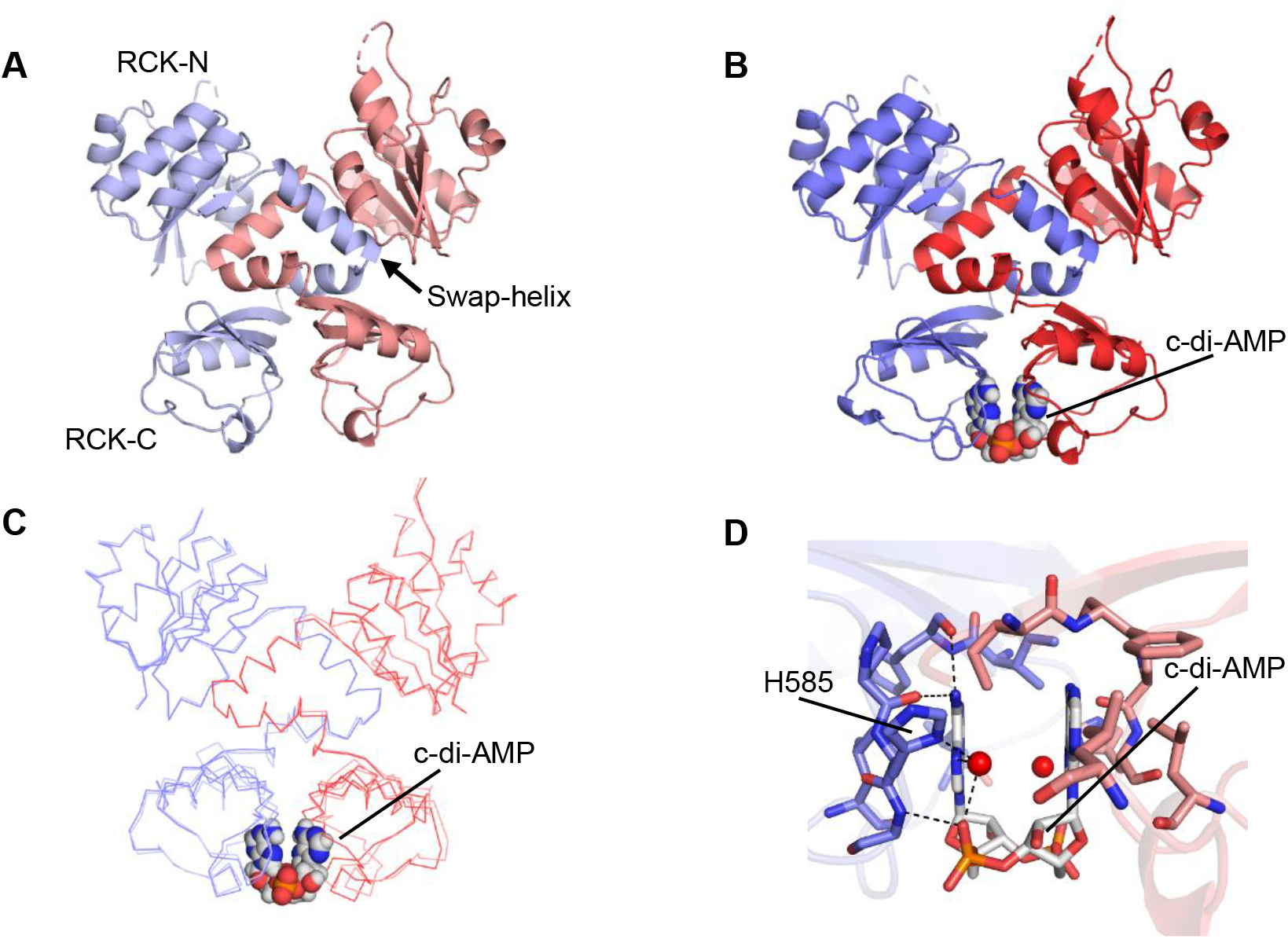
Structure of the RCK domain of CpaA with and without c-di-AMP. **A)** Structure of apo-CpaA-RCK dimer with two subunits colored in blue and red. N- and C-subdomains of RCK, as well as the swap-helix, are indicated. **B)** Structure of holo-CpaA-RCK. The two subunits of CpaA-RCK are colored with darker colors than in A). c-di-AMP is shown as spheres. **C)** Superposition of apo- and holo-CpaA- RCK, in Cα representation. **D)** Close-up view of the c-di-AMP binding site in CpaA-RCK. c-di-AMP and surrounding residues are shown as sticks. Hydrogen bonds are shown as dashed lines. Water molecules are shown as red spheres.

The CpaA-RCK dimer structure closely resembles that of the RCK dimer from the MthK K^+^ channel (24,25), where the link between the swap-helix and RCK-C is established by an α-helix and RCK-N and RCK-C subdomains sit on the same side of the dimer interface (Figure S2A). In contrast, in a RCK dimer from KtrA (17), the regulatory protein of the KtrAB cation channel complex, the linker helix is replaced by a β-strand and RCK-N and RCK-C are on opposite sides of the dimer interface (Figure S2A). Unlike what happens in the MthK and KtrAB channels, the CpaA-RCK dimer does not assemble into an octameric ring in the crystal and does not form long range helical arrangements like KtrC, another RCK regulatory protein(15). This is likely due to a reduced apolar character of the solvent exposed surface of helices αC and αD in the RCK-N subdomain of CpaA-RCK, relative to KtrA (Figure S2B). In the MthK and KtrAB octamers these helices mediate dimer-to-dimer interactions that give rise to the octameric assemblies (25,26).

The structure of the holo-protein was determined at 1.85 Å (Figure 2B and Table S2). The dinucleotide is located in the dimer interface of RCK-C, as observed before in the structures of identical subdomains from the *Staphylococcus aureus* proteins KtrA (PDB 4XTT) (22) and *S. aureus* CpaA (PDB 5F29) (23), this one a closely related orthologue of our protein (Figures S3A-B). The structure of CpaA-RCK with c-di-AMP superposes nicely with the structure of the apo-CpaA-RCK (Figure 2C), showing no significant changes in the spatial relationship between the different subdomains and just a shortening (less than 2 Å) of the distance separating residues positioned across the RCK-C interface (Figure S4). The apo- and holo-protein structures were obtained from crystals with different space-groups and cell dimensions. It is possible that a bound-like state was selected during crystallization of the apo-protein, explaining the structural similarity. We speculate that in solution the apo-protein will be less constrained, exhibiting higher flexibility at the RCK-C region, and adopting a more open conformation.

Of the interactions established by the dinucleotide with CpaA-RCK protein, it is worthwhile mentioning a water-mediated network of hydrogen bonds involving a conserved histidine (H585) (one from each subunit) and the phosphate groups and adenine bases of the dinucleotide (Figure 2D). This arrangement was previously described for *S. aureus* CpaA (Figure S3C) and is different from the interactions observed in *S. aureus* KtrA (Figure S3D), where a conserved arginine from a single-subunit slides between the two adenine bases and establishes direct and water-mediate interactions with the dinucleotide.

### c-di-AMP inactivates the CpaA cation/H^+^antiporter

To characterize the functional properties of the full-length CpaA cation/H^+^ antiporter, we first created a *cpaA* deletion mutant strain in the *Bacillus subtilis* 168 strain background and evaluated its growth phenotype in Spizizen minimal media at pH 5.0, 7.0 and 9.0, with either 2 or 100 mM KCl. We did not detect any growth differences relative to the wild-type strain, with the exception of slightly lower optical density in the stationary phase at pH 9.0 with 2 mM KCl, which disappeared with 100 mM KCl (Figure S5).

We then turned to a classical vesicle flux assay (18,27), where we prepared inside-out membrane (or everted) vesicles from the *E. coli* KNabc strain, which lacks active cation/H^+^ antiporters, expressing full-length CpaA. The vesicles were charged with protons by addition of lactate and activation of the native *E. coli* membrane bound lactate dehydrogenase (Figure 3A). CpaA activity was monitored through dequenching of the proton-sensitive dye, ACMA, after the addition of monovalent cation and exchange with protons in the vesicle lumen.

**Figure 3.**
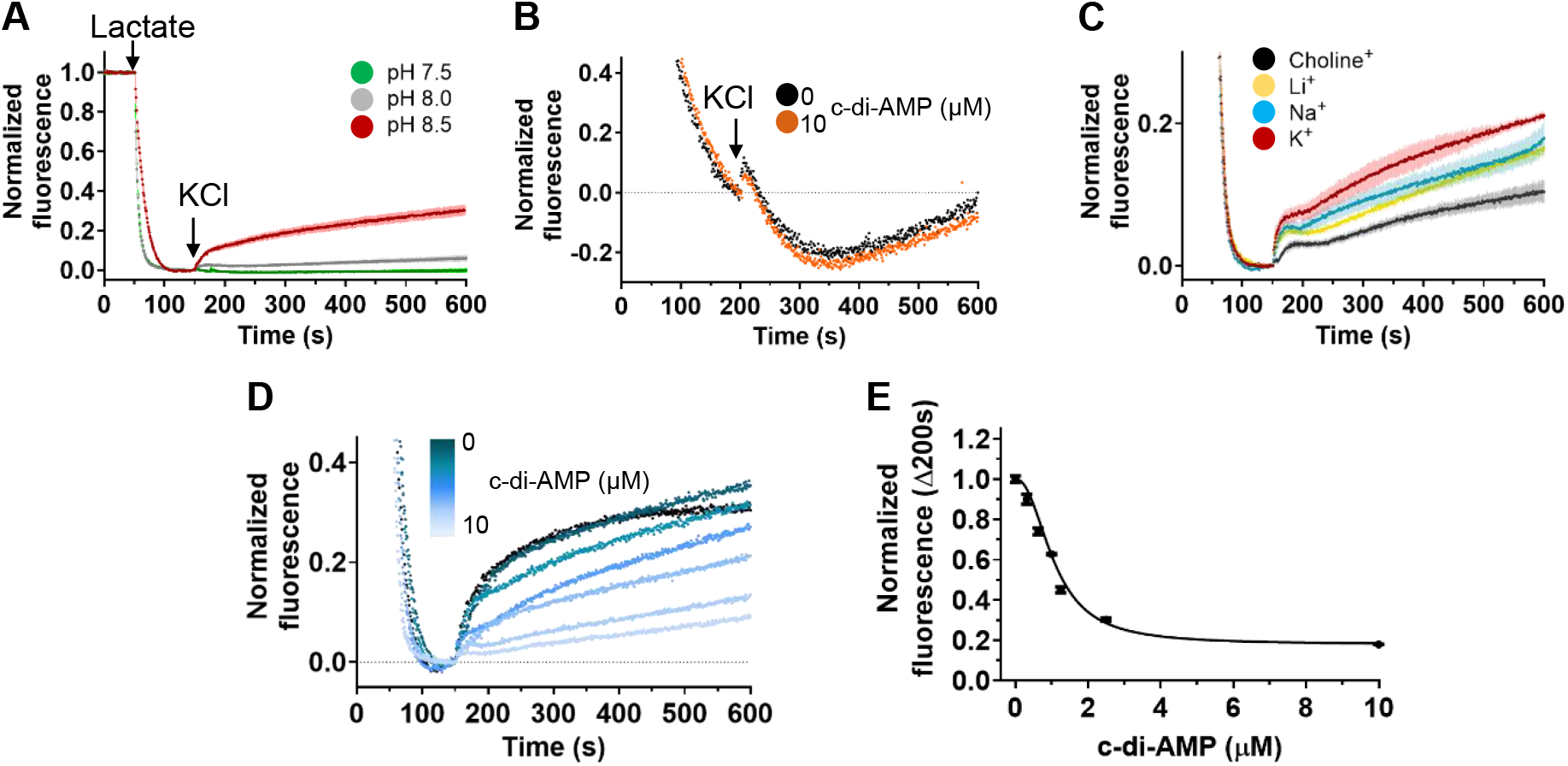
Functional characterization of *B. subtilis* CpaA. **A)** Normalized mean fluorescence curves for CpaA-containing everted vesicles obtained at pH 7.5, 8.0 and 8.5. **B)** Representative fluorescence curves for empty everted vesicles, in the absence and in the presence of c-di-AMP. **C)** Normalized mean fluorescence traces for CpaA-containing everted vesicles obtained with the addition of different monovalent cations (K^+^, Na^+^, Li^+^ and choline). **D)** Representative fluorescence curves for CpaA-containing everted vesicles, after addition of KCl, in the presence of increasing concentrations of c-di-AMP. **E)** Plot of normalized mean fluorescence determined 200 s after addition of KCl, as a function of c-di-AMP concentration. Data were fitted with Hill equation with K_1/2 inactivation_ = 1.1 ± 0.1 µM, Hill coefficient = 2.2 ± 0.2 and 0.180 for end asymptote. All monovalent cations were added at 100 mM. Mean ±SD are shown in A), C) and E) with SD shown as shaded color in A) and C) and as error bars in E).

With this assay we determined that CpaA activity is pH sensitive (Figure 3A), with higher activity at pH 8.5 relative to 8.0 and 7.5. It is also clear from these experiments that the activity level displayed by the CpaA vesicles is relatively low since fluorescence dequenching corresponds to a fraction of the lactate-induced fluorescence quenching. This is likely a reflection of low levels of protein expression and therefore of the formation of a large number of vesicles with little or no CpaA in the membrane. The low activity levels resulted in heightened sensitivity to changes in the fluorescence baseline, raising specific problems with the quantification of ion flux which are discussed below. All assays presented below were performed at pH 8.5. Importantly, vesicles prepared from cells not expressing CpaA did not show dequenching after addition of KCl (Figure 3B).

We also characterized cation selectivity by comparing the dequenching of ACMA after addition of monovalent cations K^+^, Na^+^, Li^+^ or the organic cation choline, which is too large to permeate through CpaA and functions as a negative control. As seen from the amplitude of the fluorescence dequenching, CpaA favors K^+^ over the other inorganic cations but it can also exchange Na^+^ and Li^+^ for protons (Figure 3C).

Finally, we determined the impact of c-di-AMP in the activity of CpaA. Unexpectedly, increasing concentrations of c-di-AMP reduce K^+^ flux as concluded from a decrease in the amplitude of the dequenching curves (Figure 3D). This strongly indicates that c-di-AMP inhibits CpaA activity. To quantify the c-di-AMP effect we first attempted to extract dequenching (ion flux) rate constants from exponential fits of the dequenching curves, as we had done before with KhtTU (11). However, the fits were not satisfactory due to the low activity detected. Instead, we resorted to determine the fluorescence values 200 seconds after the addition of KCl to the assay. At this time point, ACMA dequenching due CpaA activity has stabilized and the fluorescence changes are mainly due to proton leak. Normalized fluorescence was plotted as a function of c-di-AMP concentration and fitted to a Hill equation with an apparent K_1/2_ of inactivation of ∼1 µM (Figure 3E).

The unexpected inactivation by c-di-AMP in conjunction with the low activity levels in the assay made us validate these findings by generating CpaA mutants that are no longer regulated by the dinucleotide and monitor their activity in the presence of increasing ligand concentrations using the flux assay. After analysis of the c-di-AMP binding site in the CpaA-RCK structure, we selected two different residues with the view of abolishing dinucleotide binding (Figure 4). In particular, we mutated H585 to alanine, removing a side-chain that contributes to the interactions in the binding site, and G586 to serine, generating a steric clash with the ligand, and R589 to alanine, a residue far away from the binding site, as a control.

**Figure 4.**
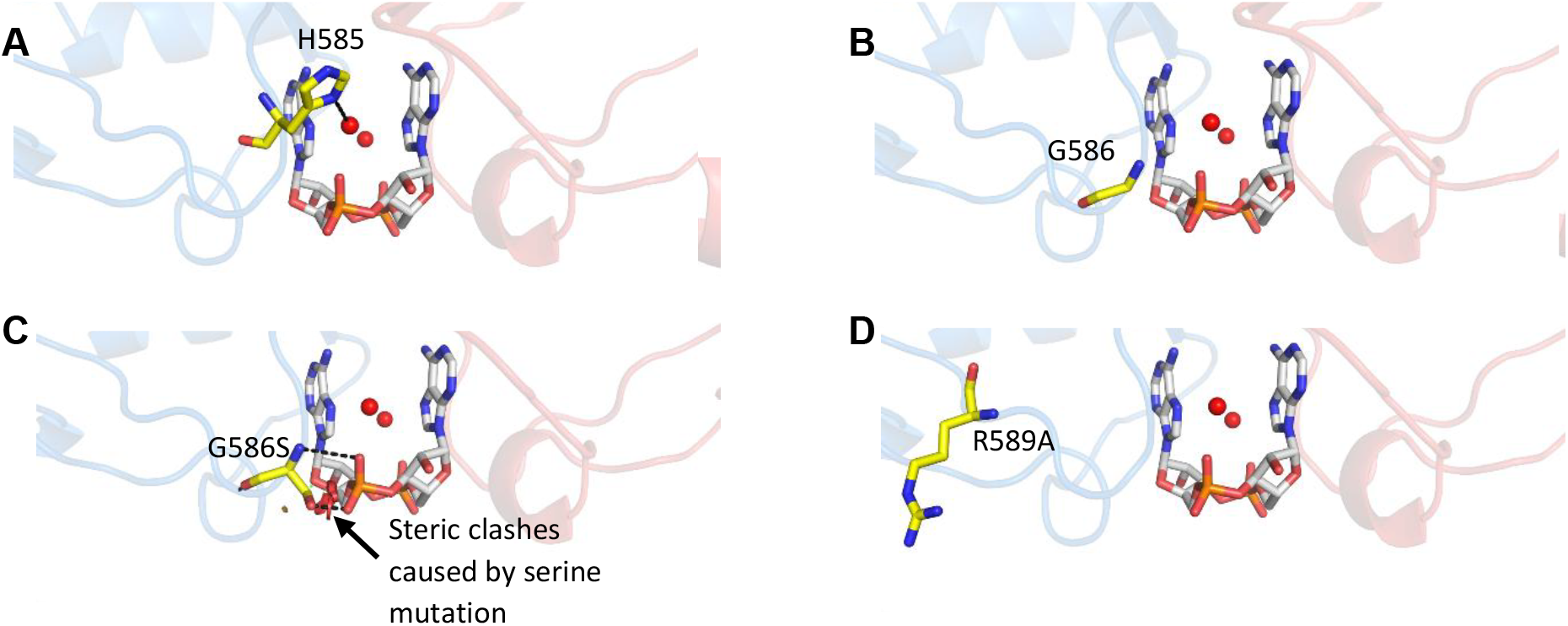
Mutated residues in the c-di-AMP binding site and vicinity. Close-up view of the c-di-AMP binding site of CpaA with c-di-AMP and selected residues represented as sticks and water molecules as red spheres. **A)** H585 interacts with c-di-AMP through a water molecule (red sphere). **B)** G586 is located on the outskirts of the ligand binding site. **C)** Modeled substitution of the G586 for a serine, highlighting the steric clashes with the phosphate group of the dinucleotide. **D)** R589 located in the vicinity of the c-di-AMP binding site is not involved in ligand interactions.

We expressed and purified these CpaA-RCK mutants as before, and together with the wild-type protein, determined their binding affinity for c-di-AMP using Isothermal Titration Calorimetry (ITC) (Figure 5). These experiments showed that wild-type protein binds c-di-AMP with a K_D_ ∼300 nM and the mutant R586A with a K_D_ = 70 nM. The difference between the K_D_ of the wild-type RCK domain and the c-di-AMP K_1/2_ of inactivation for CpaA (∼1 µM) reflects the energetic coupling between c-di-AMP binding in the RCK domain and the induction of the ligand-dependent conformational changes in the antiporter domain. In contrast, titrations with the mutants H585A and G586S showed little or no heat changes as the c-di-AMP concentration increases.

**Figure 5.**
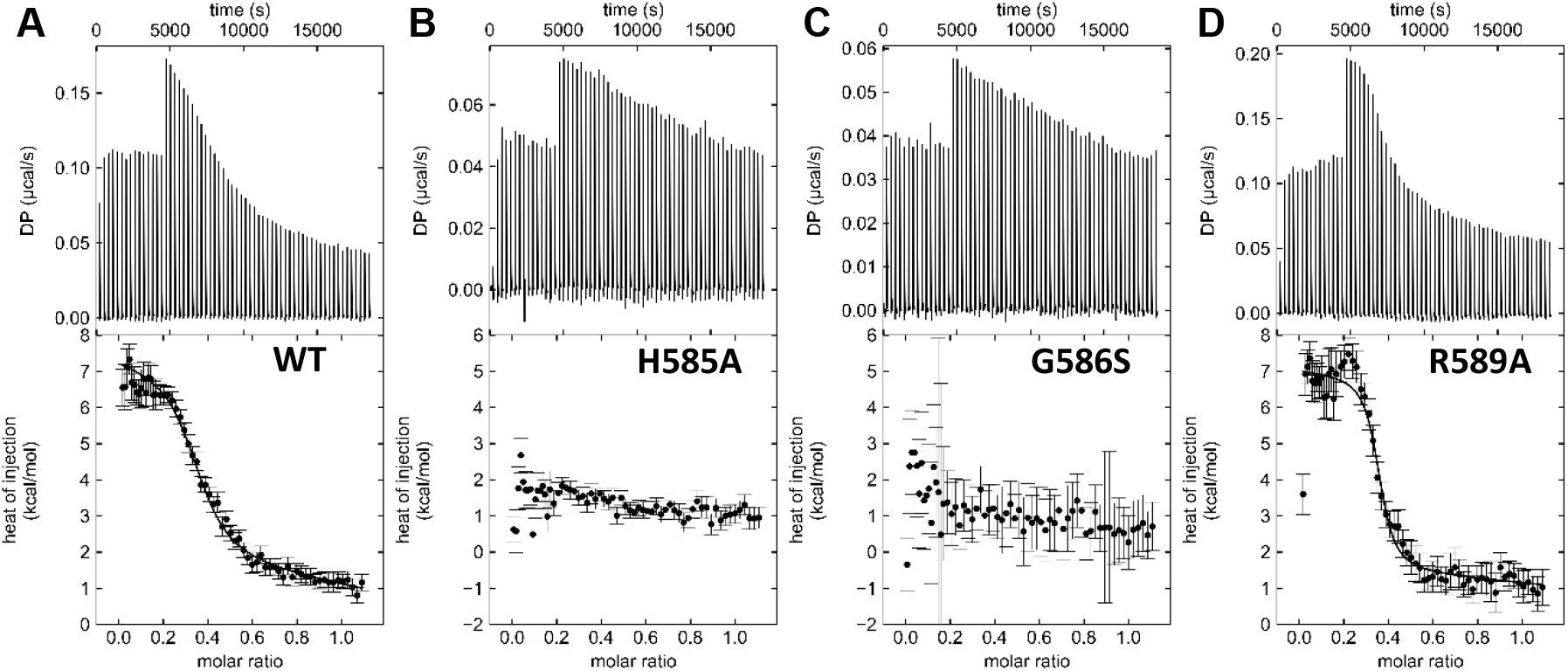
Characterization of the c-di-AMP interaction with CpaA-RCK variants. Titration of c-di-AMP into **A)** CpaA-RCK, **B)** CpaA-RCK_H585A, **C)** CpaA-RCK_G586S and **D)** CpaA-RCK_R589A. Top panels show raw titration heat values plotted as a function of time. Bottom panels show integration heat values of injectant plotted as a function of molar ratio. Curves shown in A) and D) correspond to fit of a single binding site model with n (incompetent fraction) = 0.64, ΔH = 5.2 kcal/mol, K_D_ = 0.33 µM (K_D_ 68% confidence interval: 0.19-0.56 µM) for CpaA-RCK, and n = 0.65, ΔH = 5.3 kCal/mol, K_D_ = 0.07 µM for CpaA-RCK_R589S (K_D_ 68% confidence interval: 0.01-0.22 µM).

We also analyzed the mutants using the Sypro Orange thermal shift assay (Figure S6 and Table 1). As expected, R589A displayed increased stability in the presence of 100 µM c-di-AMP (ΔTm > 6°C) that is similar to the wild-type protein (+6.1 °C). In contrast, G586S displayed no changes in thermal stability when c-di-AMP was present, indicating that the ligand does not interact with the protein. However, H585A displayed a ΔTm just above 2°C, suggesting that the mutant may bind c-di-AMP weakly, an interaction that can only be detected at the high concentration used (100 µM).

**Table 1.** Melting temperature of CpaA-RCK mutants.

| Protein variants | c-di-AMP | T<sub>m</sub> ($^\circ\text{C}$) | | | $\Delta T_m$ ($^\circ\text{C}$) |
| --- | --- | --- | --- | --- | --- |
|  |  | Replicates | Mean | SD |  |
| CpaA-RCK | - | 6 | 56.3 | 1.1 | 0 |
|  | + | 6 | 62.4 | 0.5 | +6.1 |
| CpaA-RCK H585A | - | 6 | 56.0 | 0.3 | 0 |
|  | + | 6 | 58.2 | 0.4 | +2.2 |
| CpaA-RCK G586S | - | 4 | 55.8 | 0.5 | 0 |
|  | + | 5 | 56.0 | 0.4 | +0.2 |
| CpaA-RCK R589A | - | 5 | 56.5 | 0.7 | 0 |
|  | + | 5 | 63.2 | 0.4 | +6.7 |

Temperature of melting (T<sub>m</sub>) was determined from the inflection points of fluorescence curves. $\Delta T_m$ was calculated relative to T<sub>m</sub> of the corresponding protein in the absence of ligand.

We then prepared vesicles from *E. coli* KNabc strain expressing the different CpaA mutants and performed flux assays in the same conditions as before, pH 8.5 with addition of 100 mM KCl. CpaA_R589A displayed similar sensitivity to incubation with c-di-AMP as wild-type CpaA, with a clear decrease in dequenching as ligand concentrations increase (Figure 6A).

**Figure 6.**
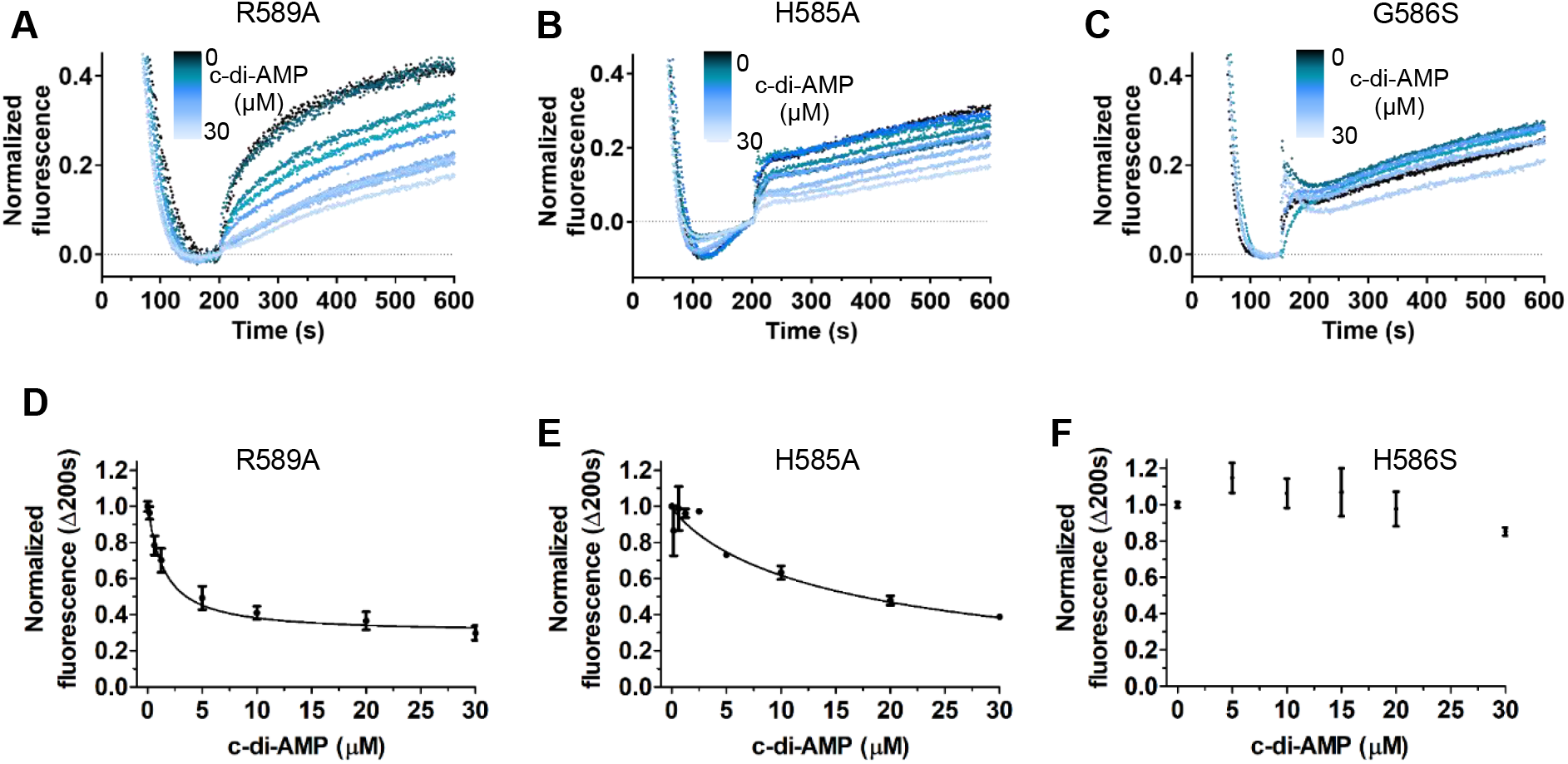
Fluorescence ion-lux assays with CpaA mutants. Representative fluorescence curves for c-di-AMP titration with **A)** CpaA_R589A, **B)** H585A and **C)** G586S. Plot of normalized fluorescence values (mean ± SD of 3 independent measurements) determined 200 s after addition of KCl, as a function of c-di-AMP concentration for **D)** CpaA_R589A, **E)** CpaA_H585A and **F)** CpaA_G586S. Data were fitted with Hill equation with K_1/2inactivation_ of 1.5 ± 0.2 µM and 10.6 ± 0.7 µM, Hill coefficient of 1.1 ± 0.1 and 1.1 ± 0.1, and end asymptote of 0.30 and 0.18 for CpaA_R589A and CpaA_H585S, respectively. KCl was added at 100 mM. Black curves correspond to assays performed without c-di-AMP.

The same was observed with CpaA_H585A but the changes in fluorescence dequenching were attenuated (Figure 6B). Finally, for CpaA_G586S, increasing c-di-AMP did not alter fluorescence dequenching relative to the assay performed in the absence of ligand (Figure 6C). Quantification of the effects confirms these conclusions. CpaA_R589A displayed a K_1/2inactivation_ of 1.5 ± 0.2 µM (Figure 6D), similar to the wild-type protein (Figure 3E), while CpaA_H585S had a K_1/2inactivation_ of 10.6 ± 0.7 µM (Figure 6E), reflecting the reduced affinity of the ligand for this mutated binding site. We could not meaningfully fit the Hill equation to the plotted data in Figure 6F, supporting the conclusion that this mutant no longer binds c-di-AMP.

Altogether, this mutant analysis confirms that the K^+^/H^+^ CpaA antiporter is inactivated by c-di-AMP.

## Discussion

Our analysis of the structure and function of the CpaA cation/proton antiporter from *B. subtilis* has confirmed that the C-terminal RCK domain contains the binding site for c-di-AMP (14) and has demonstrated that c-di-AMP regulates CpaA activity.

The structural comparison of the apo- and holo-structures of RCK revealed in detail the interactions established between the binding site residues and c-di-AMP but does not provide strong insights into the mechanism of functional regulation of the antiporter since there is only a small tightening of the dimer interface when the dinucleotide is bound.

We also showed that CpaA is a cation/H^+^ antiporter that favors K^+^ over Na^+^ but still allows sodium ion exchange. This establishes a collection of cation/proton antiporters in *Bacillus subtilis* that cover a sliding scale of cation selectivity, with KhtTU being highly selective for K^+^, CpaA with preference for K^+^ over Na^+^, NhaK with preference of Na^+^ over K^+^ and the Mrp complex displaying high selectivity for Na^+^ (11,28,29).

Importantly, we showed that c-di-AMP inactivates CpaA. This last finding was unexpected. The general view of c-di-AMP as the master regulator of the K^+^ machinery postulates that K^+^ uptake (or import) is inactivated by c-di-AMP, while K^+^ efflux (or export) is activated by this dinucleotide (10-15,20,21). Do our results mean that the CpaA K^+^/H^+^ antiporter functions as a K^+^ importer? If we assume that the transport function of this protein is reversible, then the direction of K^+^ movement will depend on the directions of the K^+^ and H^+^ concentration gradients and of the membrane electrical potential (assuming that the antiporter is electrogenic). In addition, the transport mechanism of the antiporter, through the net electrical charge (the difference between the number of protons and potassium ions for each transport cycle) moved across the membrane, also has a role in the determination of the directional movement of K^+^. A simple back of the envelope calculation, after fixing the membrane potential at-150 mV (30), shows that a K^+^/H^+^ antiporter can function as a K^+^ importer in very specific conditions, which include an external alkaline pH (higher than 7.8), so that the proton gradient is either null or directed outward, a low intracellular K^+^ concentration and a transport mechanism where the number of potassium ions transported exceeds that of protons. In particular, the intracellular K^+^ concentration would have to be lower than 50 mM at an external pH of 8.5. Intracellular K^+^ levels in bacteria are highly dynamic, reaching close to 1 M in some conditions and low tens of millimolar in others (31-33). In addition, it has been shown that c-di-AMP levels are low when intracellular K^+^ is low (3,8). It is therefore possible that CpaA contributes for the replenishment of cytosolic K^+^ but this remains to be demonstrated experimentally.

Additionally, we did not find an altered growth phenotype with a *cpaA* deletion mutant in different pH and K^+^ growth conditions. However, it has been proposed that like the Kef antiporters in *Escherichia coli*, the KhtTU K^+^/H^+^ antiporter has a role in adaptation to oxidative stress (34,35), where acidification of the cytosol contributes to the adaptation mechanism. It is not known whether c-di-AMP has a role in this mechanism but it is possible that *B. subtilis* has two K^+^/H^+^ antiporters, KhtTU and CpaA, with opposing responses to c-di-AMP to ensure that adaptation to oxidative stress involving K^+^/H^+^ antiport mechanism is available independently of the dinucleotide concentrations.

Further experiments are required to establish the physiological role CpaA and how it relates to regulation by c-di-AMP. Importantly, together with our characterization of Ktr regulation by c-di-AMP (13), which showed that the two regulatory proteins of the Ktr channels in *B. subtilis* have very distinct sensitivities to the second messenger, this study establishes that regulation of the K^+^ machinery by the c-di-AMP is more complex than previously thought.

## Experimental procedures

### Cloning and site-directed mutagenesis

The gene encoding *B. subtilis* CpaA was cloned into pBADHisB, without including the hexa-histidine tag in the construct, using primers 5’-ACCGCTCGAGAGCGCATACATCTGTCGC-3’ and 5’-GAAAACATTGGAAGGATAAGAATTCGCTT-3’. The CpaA-RCK encoding DNA fragment (residues R394 to G614) was cloned into pET15b with a hexahistidine tag at the N terminus using primers 5’-CTAGACATATGCGGGAAGAGCAGCCGGAGGAG and 5’- CTTAAGAAAACATTGGAAGGATAAGGATCCTCTAG. Mutations were generated by site-directed mutagenesis using primers 5’-GTGTGGACAGCATCGTTCCTGCAGGGGATACGAGGC and 5’-GCCTCGTATCCCCTGCAGGAACGATGCTGTCCACAC for mutation H585A, 5’- AAGTTTCAGCCTCGTATCGCTATGAGGAACGATGCTGTC and 5’-GACAGCATCGTTCCTCATAGCGATACGAGGCTGAAACTT for mutation G586S, and 5’-TTCCTCATGGGGATACGGCGCTGAAACTTGGAGACC and 5’- GGTCTCCAAGTTTCAGCGCCGTATCCCCATGAGGAA for mutation R589A. The gene fragment encoding for the DAC domain of *L. monocytogenes* CdaA from was cloned into pBAD33, using primers 5’-GCTCTAGAATCATTATTCGCTTTTGCCTC and 5’-ATATCTCCTTATTAAAGTTAAACAGGTACCCCTG.

### Expression and purification of CpaA-RCK variants

CpaA-RCK was expressed in *Escherichia coli* BL21 (DE3) grown in LB and protein expression was induced with 500 µM IPTG at OD_600 nm_ ∼0.8 at 18 °C for 18h. Protein pellets were resuspended and lysed in 50 mM Tris-HCl pH 8.0; 150 mM NaCl (lysis buffer) with protease inhibitors. Protein was first purified by affinity chromatography using a nickel-immobilized affinity column pre-equilibrated with 20 mM Imidazole in lysis buffer. After loading the column with lysis supernatant, solutions with different concentrations of Imidazole (10 mM, 20 mM, 150 mM and 300 mM) in lysis buffer were used to wash and elute protein from the beads. Eluted fractions were dialyzed overnight in lysis buffer at 4 °C in the presence of 1 μL of thrombin to cleave the hexahistidine tag. Untagged protein was separated from any un-cleaved protein by a second immobilized metal-affinity chromatography step in a column pre-equilibrated with 20 mM Imidazole 50 mM Tris-HCl; 150 mM NaCl. Protein was further purified by size-exclusion chromatography in a Superdex 200 10/300 GL column pre-equilibrated with lysis buffer. Purified protein fractions were pooled, concentrated, flash-frozen in liquid nitrogen and kept at -80 °C until needed. Protein concentration was determined by measuring absorbance at 280 nm.

### Thermal shift assay

CpaA-RCK variants and SYPRO Orange (Life Technologies) at final concentrations of 5 μM and 5x, respectively, were mixed in 50 mM Tris-HCl pH 8.0, 150 mM NaCl, 1 mM MgCl_2_ without or with 100 µM of c-di-AMP/c-di-GMP/pApA or 1 mM of CaCl_2_/ATP/ADP. Assay was performed in white 96-well PCR plates (Bio-Rad) sealed with Optical Quality Sealing Tape (Bio-Rad) with a temperature ramp from 25 to 95 ºC in 0.5 °C steps with 30 s hold time per step on a Real time PCR CFX96 and fluorescence was followed using HEX dye filter (515-535 nm excitation/560-580 nm emission). Melting curves were analyzed using the CFX Manager software (Bio-Rad) and the melting temperature was determined as the inflection point of the melting curve.

### Isothermal calorimetry titration

CpaA-RCK variants were dialyzed overnight against 50 mM HEPES pH 8.0, 120 mM KCl, 30 mM NaCl, 1 mM MgCl_2_, 1 mM TCEP (ITC dialysis buffer), with repeated buffer exchanges. Titrations were performed in a MicroCal VP-ITC instrument (GE Healthcare) at 15 °C with 20 μM of protein in the cell titrated with 100 μM c-di-AMP (in ITC dialysis buffer). Three titration experiments were performed for each protein and globally fit with a single-binding site model in SEDPHAT (36) after pre-processing with NITPIC (37).

### Crystallization and structure determination

Crystals of apo CpaA-RCK were obtained at 20ºC from an 8 mg/ml protein solution (in 20 mM Tris-HCl pH 8.0, 150 mM NaCl) and a Morpheus crystallization screen solution composed of 0.3 M magnesium chloride hexahydrate, 0.3 M calcium chloride dihydrate, 0.1 M buffer system pH 7.5 (1.0 M Sodium HEPES, MOPS (acid)), 30% v/v precipitant mix (40% v/v PEG 500 MME; 20% w/v PEG 2,000). Crystals were flash-frozen directly in liquid nitrogen. Crystals of holo CpaA-RCK were obtained at 20 ºC in presence of c-di-AMP from an 8 mg/ml protein solution (in 20 mM Tris-HCl pH 8.0; 150 mM NaCl, 1 mM c-di-AMP) and a crystallization solution composed of 28% PEG 8,000, 0.1 M MES pH 6.5, 0.3 M MgCl_2_. Crystals were cryoprotected with 30.8% PEG 8,000, 0.110 M MES pH 6.5, 0.330 M MgCl_2_, 15% (v/v) glycerol, 0.300 uM c-di-AMP before being flash-frozen in liquid nitrogen.

### Data collection and processing

Diffraction data were collected from cryo-cooled (100 K) single crystals at ALBA (Barcelona, Spain) and the French National Synchrotron Source (SOLEIL Gif-sur-Yvette, France) synchrotrons. Datasets were processed with MOSFLM. X-ray diffraction data collection and processing statistics are summarized in Table S2.

### Structure determination, model building and refinement

Structure of CpaA-RCK with c-di-AMP was solved by molecular replacement with Phaser and CCP4 (38,39) using the coordinates of the C-terminal subdomain of the RCK domain of *S. aureus* CpaA (PDB 5F29) and of the N-terminal subdomain of *B. subtilis* KtrA from (PDB 4J91) as search models. The structure of apo CpaA-RCK was solved by molecular replacement using the structure of CpaA-RCK with c-di-AMP as search model. Alternating cycles of model building with Coot (40) and refinement with PHENIX (41) were performed until model completion. Refinement statistics are summarized in Table S2.

### Preparation of everted membrane vesicles

Everted-vesicles prepared from KNabc *E. coli* strain expressing CpaA alone showed little or no antiport activity. Knowing that c-di-AMP was able to bind CpaA-RCK, we co-expressed CpaA with the DAC domain of *L. monocytogenes*, giving the *E. coli* cells the ability to produce c-di-AMP and increasing the antiport activity in vesicles. Therefore, KNabc cells were transformed with pBAD33-DAC together with pBADHisB-*cpaA* (for CpaA), pBAD-*cpaA*H585A (for CpaA-H585A), pBADHisB-*cpaA*G586S (for CpaA-G586S), pBADHisB-*cpaA*R589A (for CpaA-R589A) or pBADHisB (for empty everted vesicles), and were grown at 37 °C in LBK to OD_600 nm_ ∼0.8. Protein expression was induced with 0.002% arabinose for 3 h at 37 °C.

Cells were harvested and everted membrane vesicles were prepared in 20 mM HEPES pH 7.4, 140 mM choline chloride, 250 mM sucrose, 0.5 mM DTT (vesicle buffer). Briefly, cells were washed once in vesicle buffer and lysed by a single passage through a French press cell at 4,000 psi. Lysate was then incubated at room temperature with 10 mM MgCl_2_ and 20 µg/mL DNAse. After removal of unbroken cells by a 5,000xg centrifugation at 4 °C for 20 min, everted vesicles were collected by ultracentrifugation at 100,000xg and 4 °C for 1 h. Everted vesicles were washed twice with vesicle buffer by repeating resuspension and ultracentrifugation. Washed vesicles were resuspended in 1 mL of vesicle buffer per gram of original wet cell weight, flash-frozen in liquid nitrogen and kept at -80 °C until needed. For each everted vesicle preparation, the total protein concentration was determined with Bradford reagent and a calibration curve determined with Bovine Serum Albumin as a standard.

### Antiport Assays

Fluorescence-based flux assays were performed as before (11) with the following modifications. Everted vesicles were diluted in 15 mM Tris-HCl pH 8.5 (or pH 7.5 or pH 8.0); 140 mM Choline Chloride; 5 mM MgCl_2_ (vesicle assay buffer) and incubated during 3 minutes in the presence of 2 µM ACMA, added from a 300 µM stock prepared in DMSO. Assays were started by adding 4 mM lactic acid from a 600 mM stock solution diluted in Tris (1M) to promote a respiration-generated ΔpH. Proton efflux was then induced by adding 100 mM KCl for assays at pH 7.5, pH 8.0 and for most assays at pH 8.5, including all c-di-AMP titration assays; for cation selectivity assays, proton efflux was induced by adding 100 mM of KCl, NaCl, LiCl or choline chloride. For c-di-AMP titrations, nucleotide was added to vesicles together with ACMA, to a final concentration that varied between 156 nM to 30 µM in the case of vesicles with mutant variants and between 3 nM to 10 µM in the case of vesicles with wildtype protein. All assays with everted vesicles experiments were performed at least in triplicate using a single batch of everted vesicles. Fluorescence raw data was normalized as a fraction of fluorescence before addition of lactate. For c-di-AMP titrations, fluorescence values measured 200 s after addition of KCl were normalized to the fluorescence for curves recorded in the absence of dinucleotide. Values were plotted as a function of c-di-AMP concentration and fitted with a Hill equation

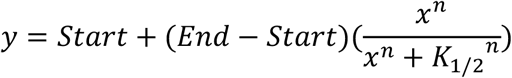

where *K*_*1/2*_ is the constant of inactivation at 50%, *n* is the Hill coefficient, *Start* was fixed as 1 and *End* is the bottom asymptote.

### *B. subtilis* culture conditions

*B. subtilis* strains 168 and 168Δ*cpaA* were grown in Spizizen Minimal Medium (SMM) (13) with variations. Briefly, 5 mL pre-cultures were grown in SMM with 2 mM KCl at 37 °C and with 220 rpm agitation for ∼5 h. Cells were washed with 150 mM NaCl and then inoculated to a final OD_600nm_ of ∼0.2 into a 96-well plate in SMM medium at different pH (5.0, 7.0 and 9.0) and at different K^+^ concentration (2 mM and 100 mM KCl). SMM media were initially prepared as a 5x concentrated media without KCl and phosphate. The pH was then adjusted with 100 mM MES pH 5.0, 100 mM Bis-Tris Propane and 100 mM Bis-Tris Propane pH 9.0 and KCl adjusted from a 3 M stock. The plate was incubated at 37 °C for ∼18 h with 600 rpm agitation. Cell growth was monitored by measuring the OD_600nm_ every 15 minutes in a CLARIOstar Multi-detection microplate reader.

## This article contains supporting information

## Acknowledgments

We acknowledge the SOLEIL and ALBA synchrotrons and the scientific platforms of i3S (Biochemical and Biophysical Technologies and X-ray Crystallography) for access and thank their staff for help with data collection.

## Funding and additional information

This work was supported by Fundação para a Ciência e a Tecnologia (FCT), under the ERC-Portugal program, through the project ‘The cellular organization and molecular function of the K^+^ machinery in a bacterium’ and by “La Caixa” Foundation under the project code LCF/PR/HR22/52420004.

