## Supplemental information for "c-di-AMP inactivates a K^+^/H^+^ antiporter in *Bacillus subtilis*"

**Table S1. Melting temperature of RCK domain**

| Ligand | Tm (°C) | | | ΔTm (°C) |
|---|---|---|---|---|
| | N | Mean | SD | |
| RCK (alone as control) | 6 | 56.5 | 0.6 | - |
| + $Ca^{2+}$ | 3 | 57.5 | 0.0 | +1.0 |
| + c-di-AMP | 3 | 63.0 | 0.0 | +6.5 |
| + c-di-GMP | 3 | 56.5 | 0.0 | 0 |
| + pApA | 5 | 56.7 | 0.3 | +0.2 |
| + ATP | 3 | 56.3 | 0.4 | -0.2 |
| + ADP | 3 | 56.5 | 0.0 | 0 |

Temperature of melting (Tm) was determined from the inflection point of melting curves. ΔTm was calculated relative to Tm of protein alone. N is the number of measurements.

**Table S2. X-ray diffraction data and refinement statistics.**

| | CpaA-RCK with c-di-AMP | CpaA-RCK |
|---|---|---|
| **Data** | | |
| Unit cell dimensions | a=110.6 b=120.1 c=36.1 (Å) α=90 β=90 γ=90 (º) | a=36.2 b=93.0 c=25.3 (Å) α=90 β=90 γ=90 (º) |
| Space group | $P2\ 2_1\ 2_1$ | $P2_1\ 2_1\ 2_1$ |
| Resolution limit (Å) | 40.7 -1.85 (1.89 -1.85) | 46.5-2.19 (2.26-2.19) |
| $R_{merge}$ | 0.086 (2.873) | 0.113 (1.674) |
| $R_{meas}$ | 0.089 (3.013) | 0.115 (1.707) |
| Number of observations | 560,130 (27208) | 598,292 (48862) |
| Number of unique observations | 42,129 (2504) | 22,502 (1889) |
| Mean I/(σI) | 15.5 (0.8) | 21.97 (2.58) |
| $CC_{1/2}$ | 1.00 (0.442) | 1.00 (0.802) |
| Completeness (%) | 99.8 (97.1) | 99.6 (96.2) |
| Multiplicity | 13.3 (10.9) | 26.6 (25.9) |
| **Refinement** | | |
| Resolution range (Å) | 40.7 - 1.85 | 46.5 - 2.19 |
| $R_{work}$ (%) | 20.25 | 21.80 |
| $R_{free}$ (%) | 24.22 | 25.56 |
| Number of residues | 416 | 416 |
| Number of waters | 70 | 23 |
| Rmsd bond length (Å) | 0.013 | 0.005 |
| Rmsd angles (º) | 1.55 | 0.88 |
| Ramachandran favoured (%) | 98 | 98 |
| Ramanchandran outliers (%) | 0 | 0 |

Values in parenthesis are for high-resolution shell.

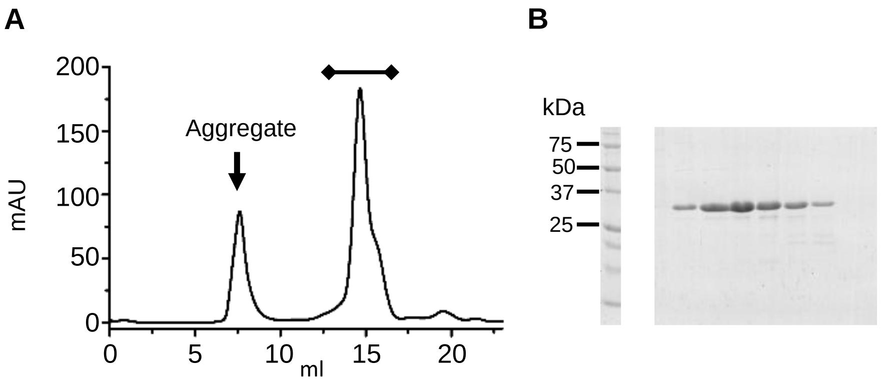


**Figure S1: Purification of the RCK domain of CpaA. A)** Size-exclusion chromatogram of the CpaA-RCK. Peak at ~7.5 ml corresponds to aggregated protein, while peak at ~15 ml corresponds to CpaA-RCK protein. **B)** SDS-PAGE of the size-exclusion fractions spanning peak at 15 ml.

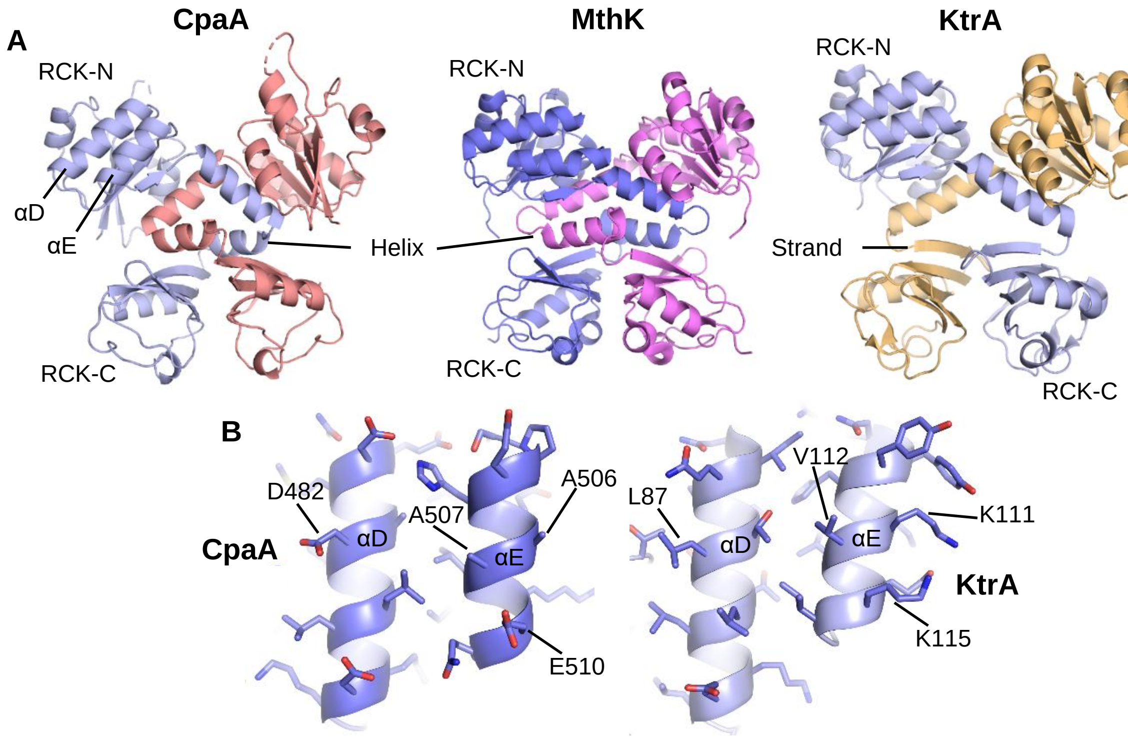


**Figure S2: Structure of the RCK domains of CpaA, MthK and KtrA. A)** RCK dimers of *B. subtilis* CpaA cation/$H^+$ antiporter, *Methanothermobacter thermautotrophicus* MthK $K^+$ channel (PDB 1LNQ) and *B. subtilis* KtrA regulatory protein (PDB 4J90) are shown as cartoon. The two subunits of each dimer are colored blue and red (for CpaA) or pink (for MthK) or yellow (for KtrA). RCK-N and -C subdomains are indicated as are the α-helices and β-strand that link the RCK-C subdomain. Helices αD and αE are indicated for CpaA. **B)** Solvent exposed surface of helices αD and αE in CpaA-RCK (left) and KtrA (right) are shown as cartoon with residues represented as sticks. The differences in the chemical character of these surfaces include the substitution of an aspartate (D482) in αD of CpaA for a leucine (L87) in KtrA, while in αE, two alanines (A506 and A507), with their short side chains, are replaced by a lysine (K111), with its long aliphatic side chain, and a valine (V112). Additionally, a glutamate (E510) in CpaA is replaced in KtrA for a lysine (K115). These differences appear sufficient to reduce in CpaA the hydrophobic effect that mediates the interaction between RCK dimer in KtrA.

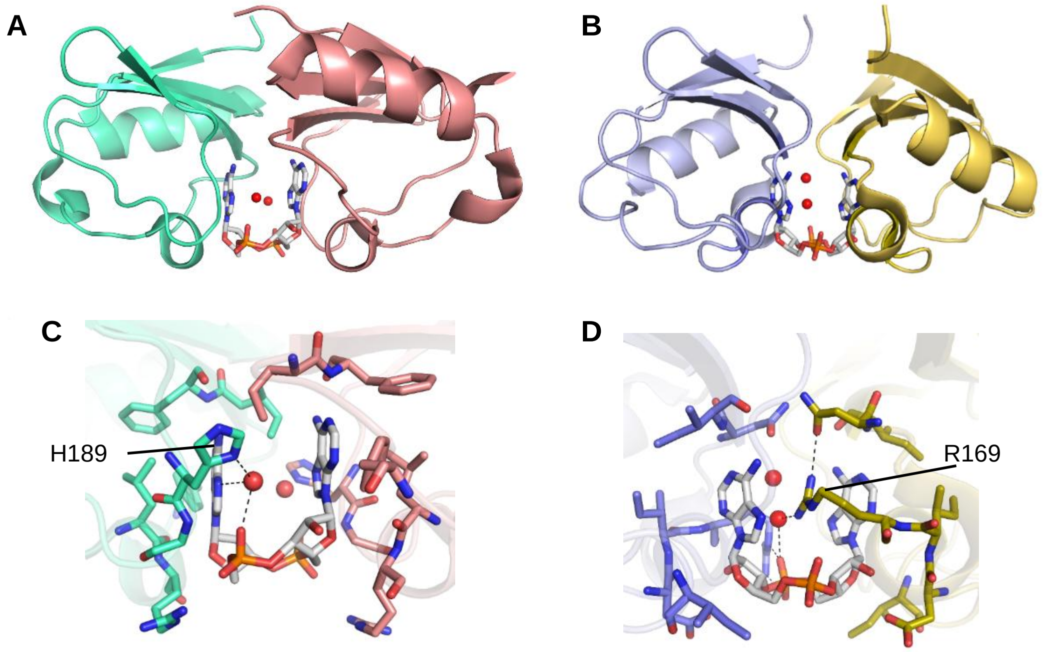


**Figure S3: RCK-C subdomains of CpaA and KtrC. A)** RCK-C subdomains of *S. aureus* CpaA cation/$H^+$ antiporter (PDB 5F29) represented as cartoon, with c-di-AMP, shown as sticks, located between the C-lobes**.** **B)** *S. aureus* KtrA regulatory protein (PDB 4XTT) shown as in A). **C)** Close-up view of the c-di-AMP binding site of *S. aureus* CpaA. c-di-AMP and the residues surrounding the ligand are shown as sticks. Water molecules involved in ligand binding are shown as red spheres. Hydrogen-bonds are shown as dashed lines. **D)** Close-up view of the c-di-AMP binding site of *S.aureus* KtrA. c-di-AMP, surrounding residues and water molecules are shown as in C.

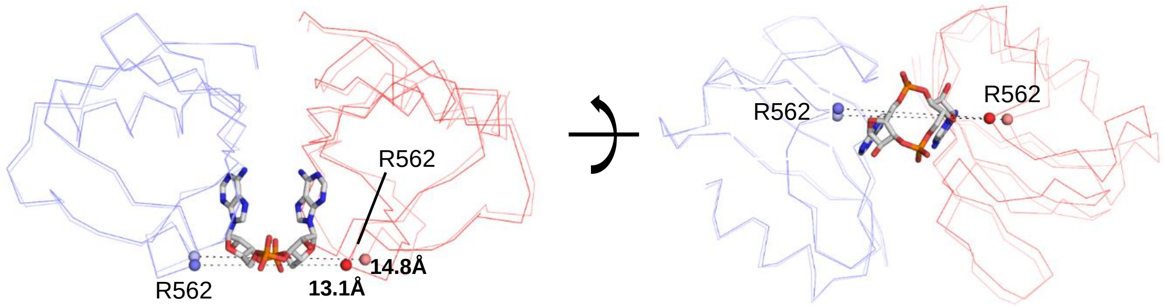


**Figure S4: Superposition of RCK-C subdomain from CpaA with and without c-di-AMP.** Structure of RCK-C of CpaA with (intense colors) and without (lighter colors) c-di-AMP are superposed through the blue subunit, are shown as Cα trace. c-di-AMP is shown as stick. Spheres indicate Cα position of R562 in the subunits (red and blue) of both structures (light and dark colors). Distances indicated separate R562 across the interface in the c-di-AMP bound (13.1Å) and unbound (14.8Å) structures.

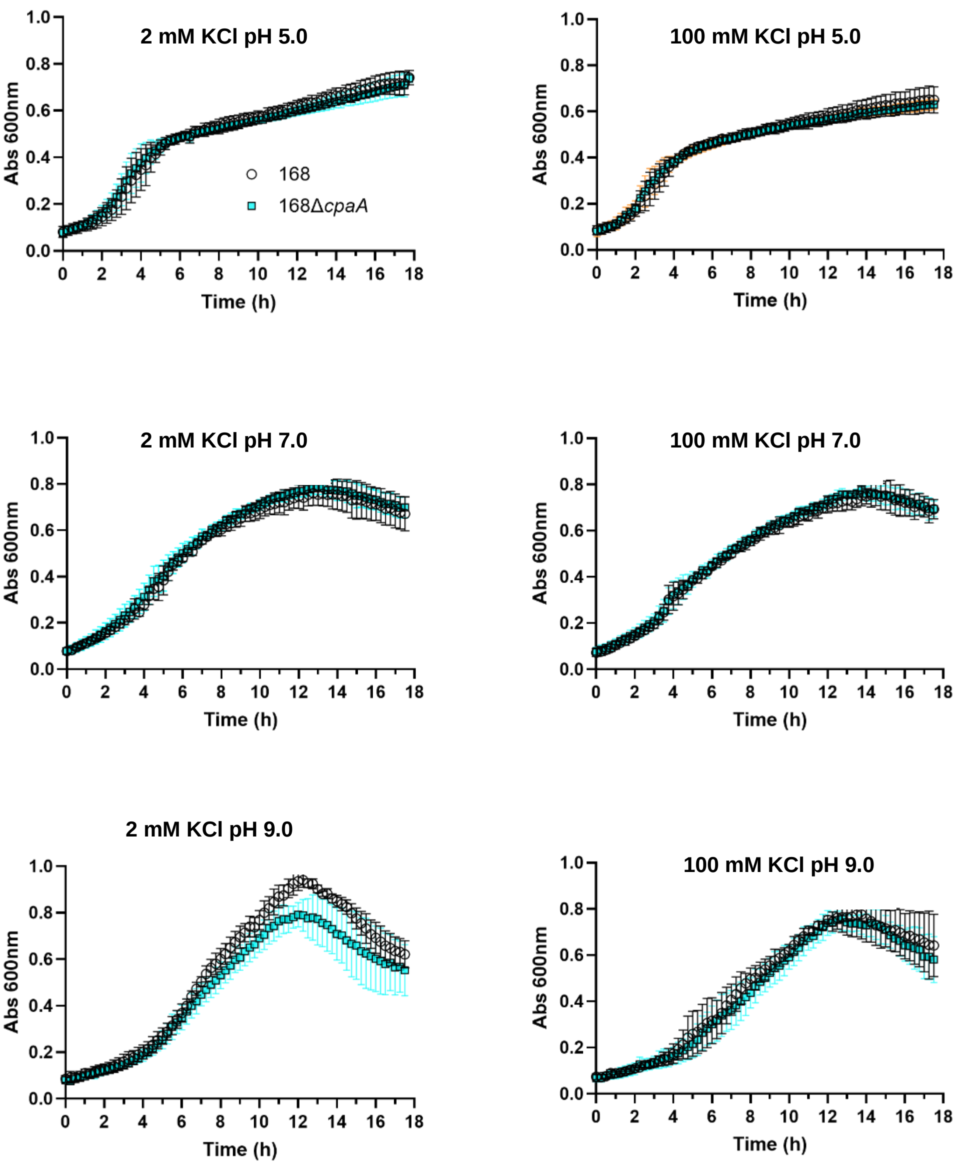


**Figure S5: Growth phenotypes of *B. subtilis cpaA* deletion mutant.** Wild-type (168, empty circles) and mutant (168Δ*cpaA*, cyan circles) strains were grown in SMM media buffered at pH 5.0 (top), 7.0 (middle) or 9.0 (bottom) with 2 (left) or 100 mM KCl(right).

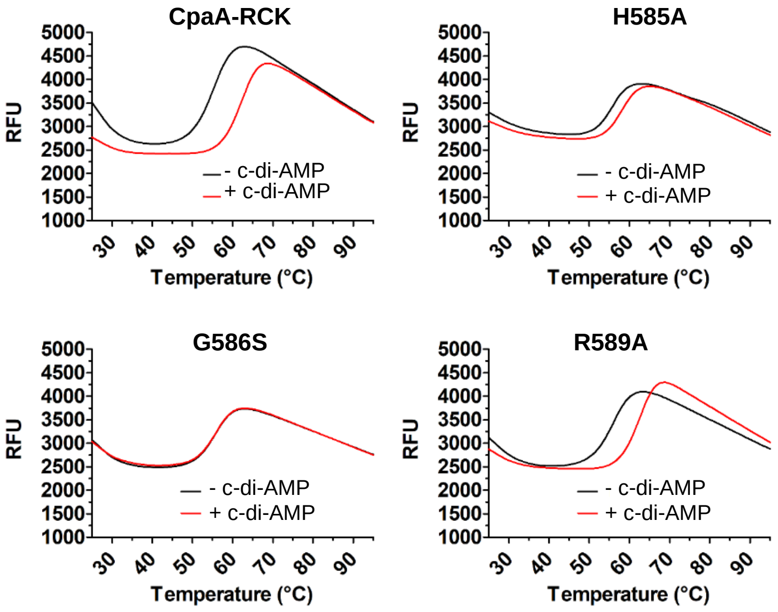


**Figure S6: Thermal shift assay of CpaA-RCK variants.** Representative fluorescence curves of Sypro Orange as a function of temperature for the wild-type CpaA-RCK (top, left) and CpaA-RCK mutants CpaA-RCK_H585A (top right), CpaA-RCK_G586S (bottom left) and CpaA-RCK_R589A (bottom right), in the absence and presence of 100 µM c-di-AMP.
